## Supplemental figures for "B cell receptor dependent enhancement of dengue virus infection"

### Supplemental Figure 1

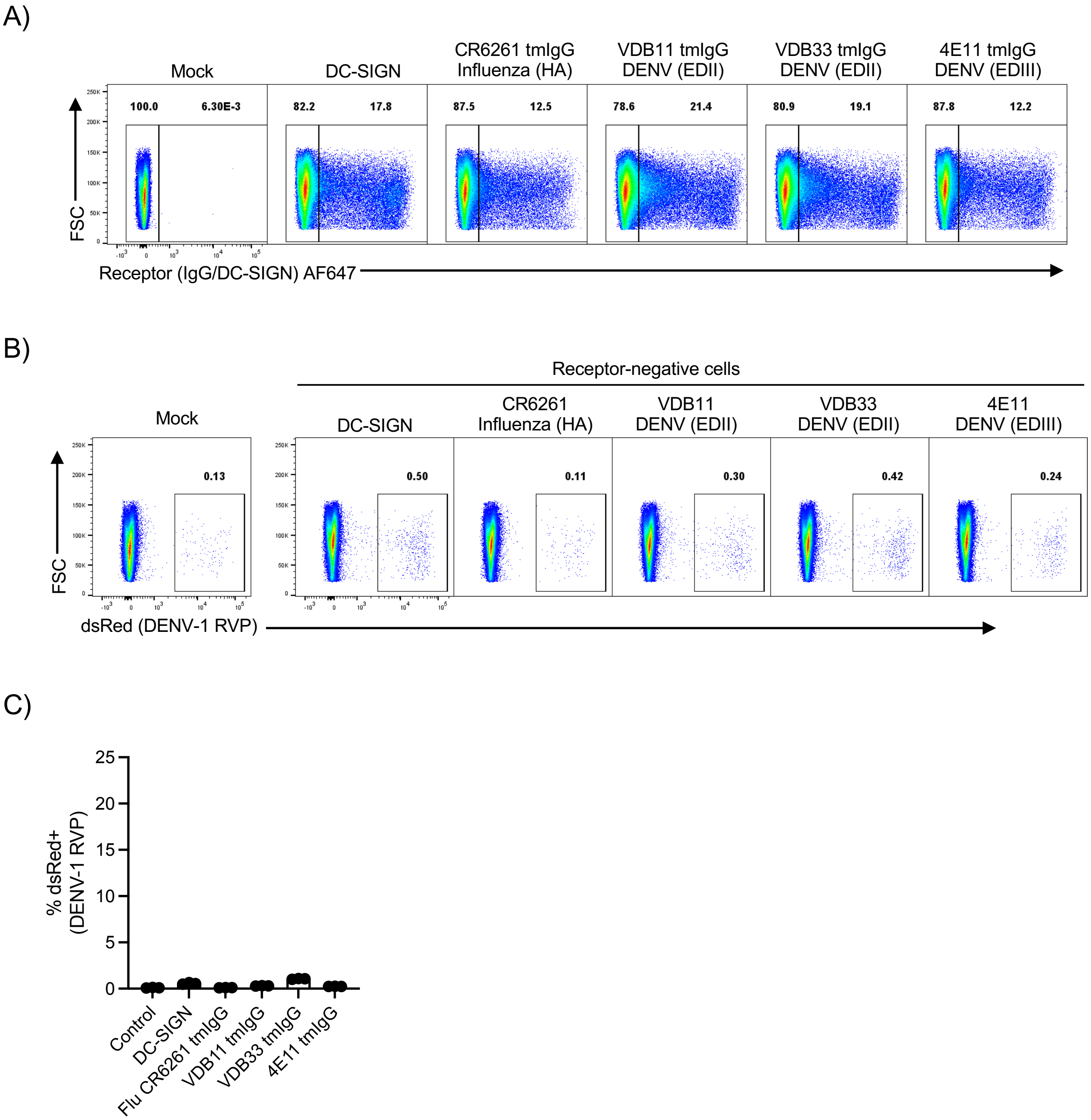


**Supplemental Figure 1. A)** Expression and gating of tmIgG and DC-SIGN in transfected 293T cells **B)** Representative flow cytometry plots showing the frequency of DENV-1 RVP infected cells within the receptor-negative gate of tmIgG and DC-SIGN in transfected 293T cells 24hrs after RVP exposure **C)** Quantification of DENV-1 RVP infected cells within the receptor-negative gate of tmIgG and DC-SIGN in transfected 293T cells 24hrs after RVP exposure

### Supplemental Figure 2

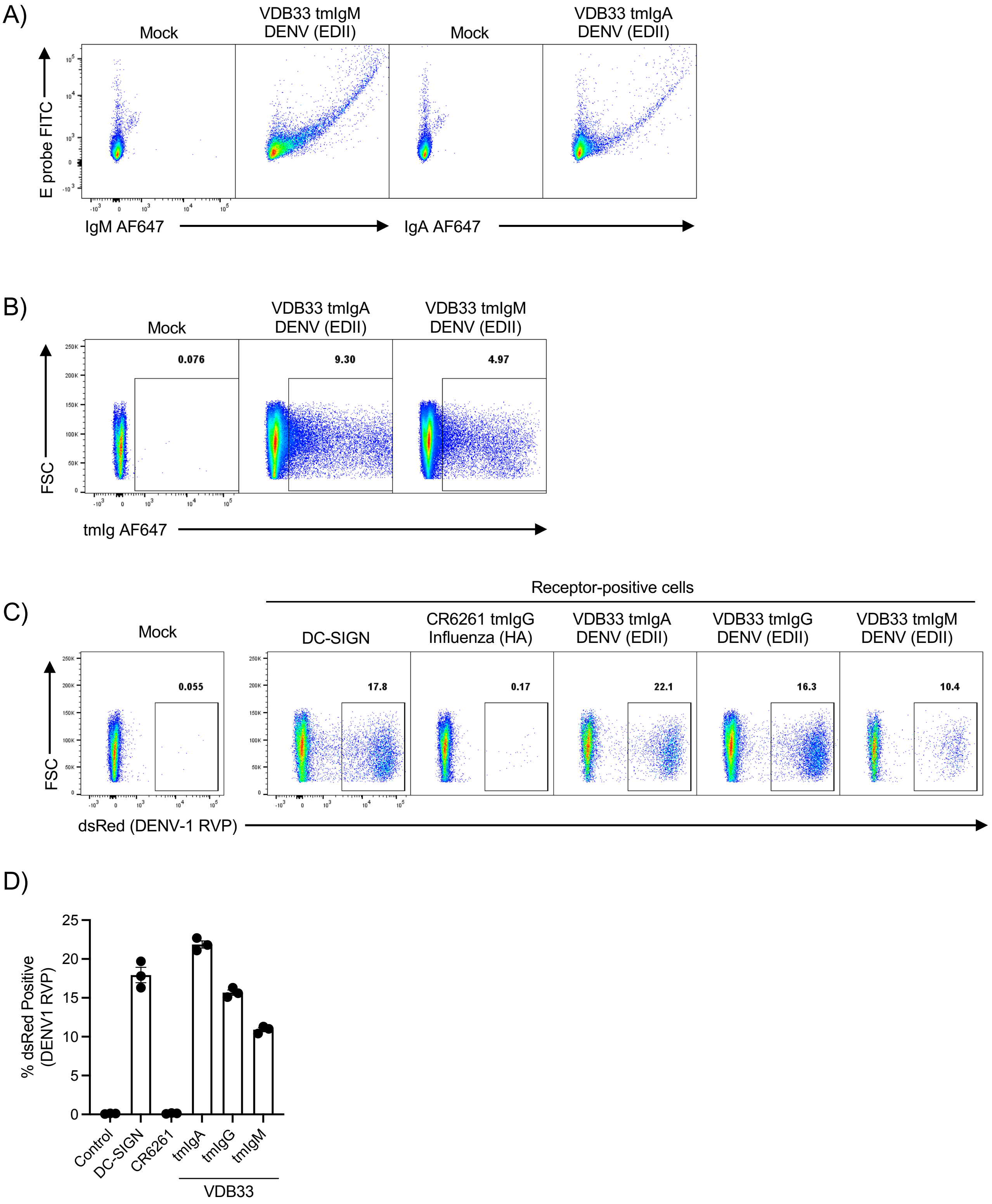


**Supplemental Figure 2. A)** Expression and DENV E protein binding activity of VDB33 tmIgM and tmIgA expression constructs. **B)** Expression and gating of tmIgM and tmIgA expression in transiently-transfected 293T cells **C)** Representative flow cytometry plots showing the frequency of DENV-1 RVP infected cells within the DC-SIGN, CR261 tmIgG, VDB33 tmIgA, VDB33 tmIgG, and VDB33 tmIgG positive 293T cells 24 hours after infection **C)** Quantification of DENV-1 RVP infected cells within the DC-SIGN, CR261 tmIgG, VDB33 tmIgA, VDB33 tmIgG, and VDB33 tmIgG positive 293T cells 24 hours after infection

### Supplemental Figure 3

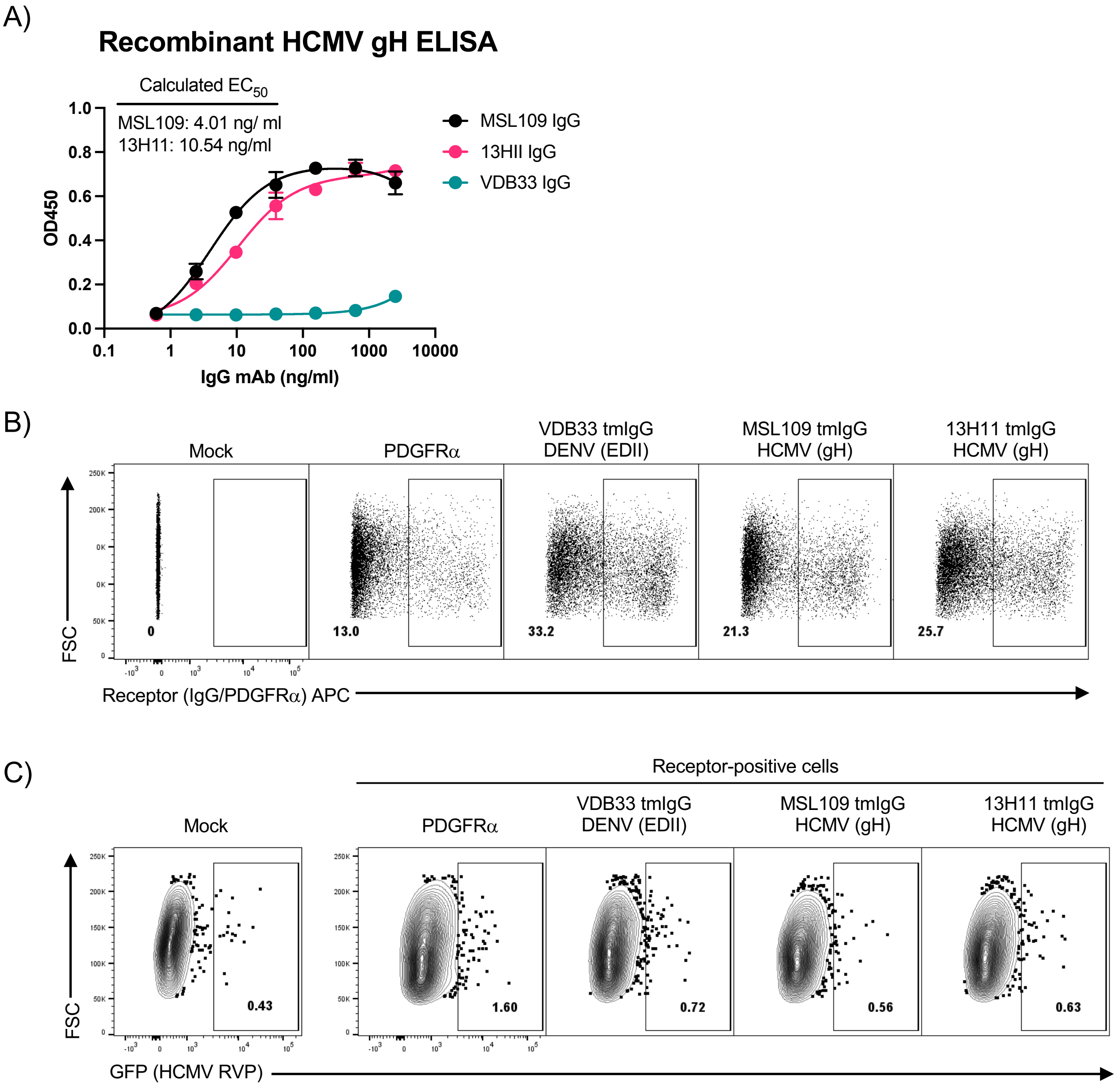


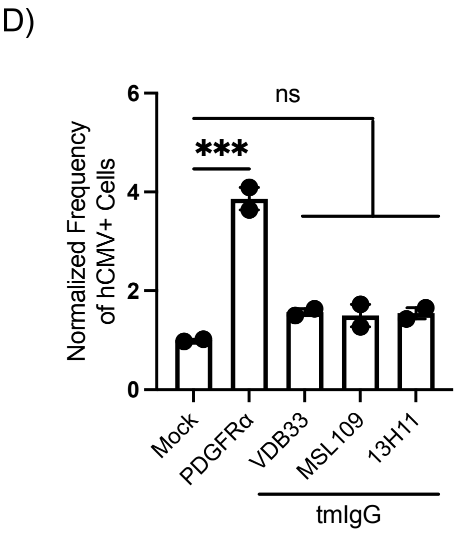


**Supplemental Figure 3. A)** Binding of the indicated IgG isotype mAbs to recombinant/purified HCMV gH protein as quantified by ELISA. **B)** Expression and gating of tmIgG and PDGFRα expression in transiently-transfected 293T cells **C)** Representative flow cytometry plots showing the frequency of HCMV SV40-GFP infected cells within the receptor-positive gate of tmIgG and PDGFRα transfected 293T cells 24hrs after virus inoculation. Cells infected at an MOI of 1 **D)** Quantification of HCMV SV40-GFP infected cells within the receptor-negative gate of tmIgG and PDGFRα in transfected 293T cells 24hrs after virus inoculation

#### Supplemental Figure 4

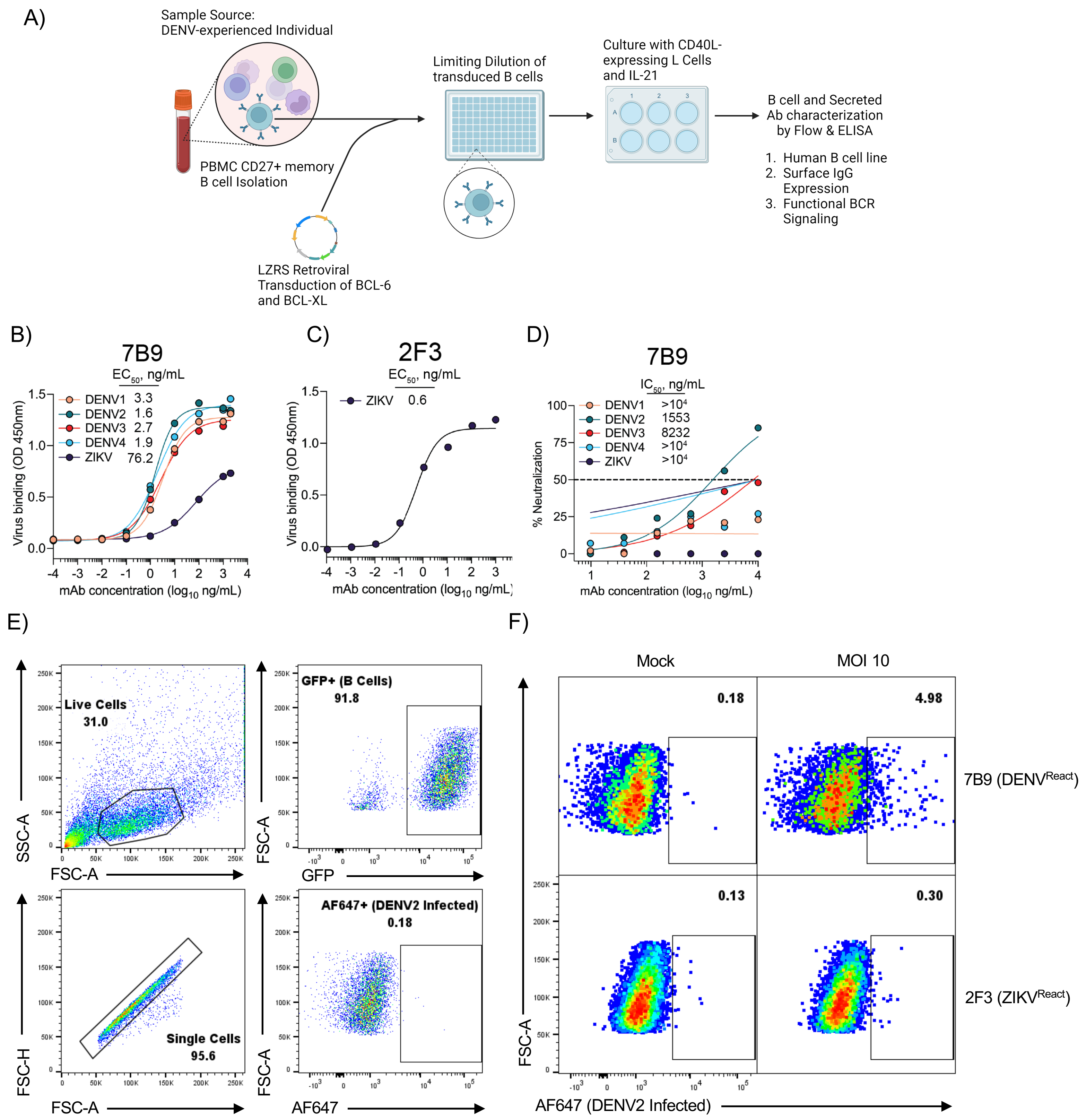


**Supplemental Figure 4. A)** Schematic representation of 7B9 and 2F3 cell line generation and maintenance. **B)** Virus binding ELISA data for mAbs expressed by 7B9 and C) 2F3 cell lines. **D)** DENV and ZIKV neutralization profiles of mAb expressed by cell line 7B9. **E)** Gating scheme for flow cytometry analysis of DENV-infected 7B9 and 2F3 cell lines. **F)** Representative flow cytometry plots showing the frequency of DENV-infected 7B9 and 2F3 cell lines after DENV-2 exposure.

### Supplemental Figure 5

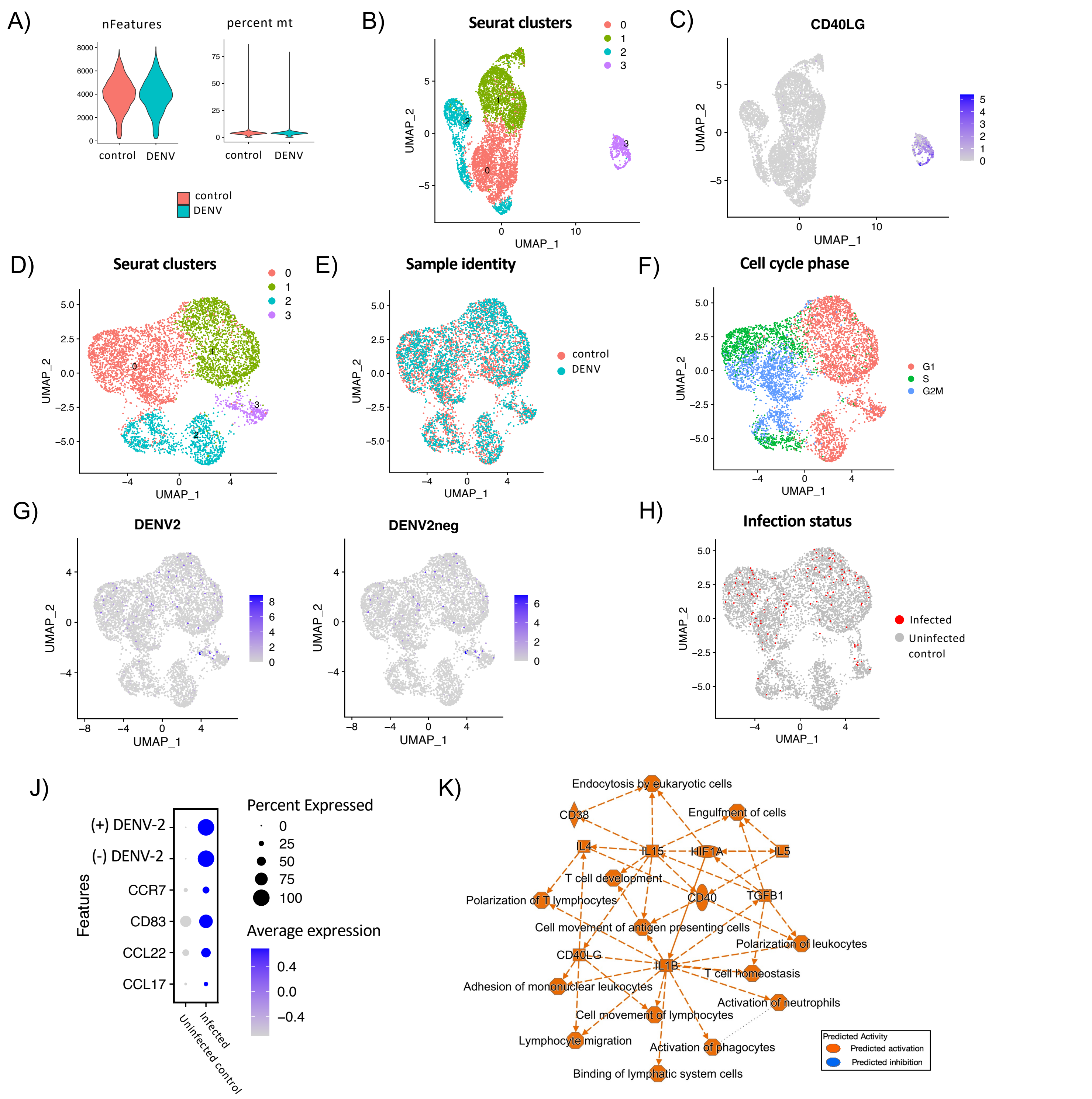


**Supplemental Figure 5. Quality control metrics and integrated UMAP projections of scRNA seq data**. **(A)** Violin plots indicating number of features (nFeatures) and percentage mitochondrial content (percent mt) pre-filtering of scRNAseq data. **(B)** Integrated UMAP projection of cells derived from all conditions prior to subsetting for removal of CD40 ligand positive feeder cell population, and **(C)** Feature plot indicating CD40 ligand (CD40LG) expression, highlighting the feeder cell population (cluster 3) which was removed for subsequent analysis. **(D)** UMAP projections of scRNAseq data indicating Seurat clusters, **(E)** Sample origin, and **(F)** Cell cycle phase. (**G**) Feature plot indicating DENV (+) and (-) sense RNA expression. **(H)** Imputed cell labelling of infected (positive for both DENV positive and negative sense RNA) and uninfected control/bystander cell populations. (G) Dot plot highlighting selectively upregulated and downregulated DEGs in infected (expressing both DENV (+) and (-) RNA) compared to uninfected control cells (lacking both DENV (+) and (-) RNA expression). Average expression and percent of cells expressing a given transcript are indicated. (H) Ingenuity Pathway Analysis (IPA) Graphical summary of predicted pathway enrichment in infected cells compared to uninfected controls based on differential gene expression between infected (expressing both DENV (+) and (-) RNA) compared to uninfected control cells (lacking both DENV (+) and (-) RNA expression).

### Supplemental Figure 6

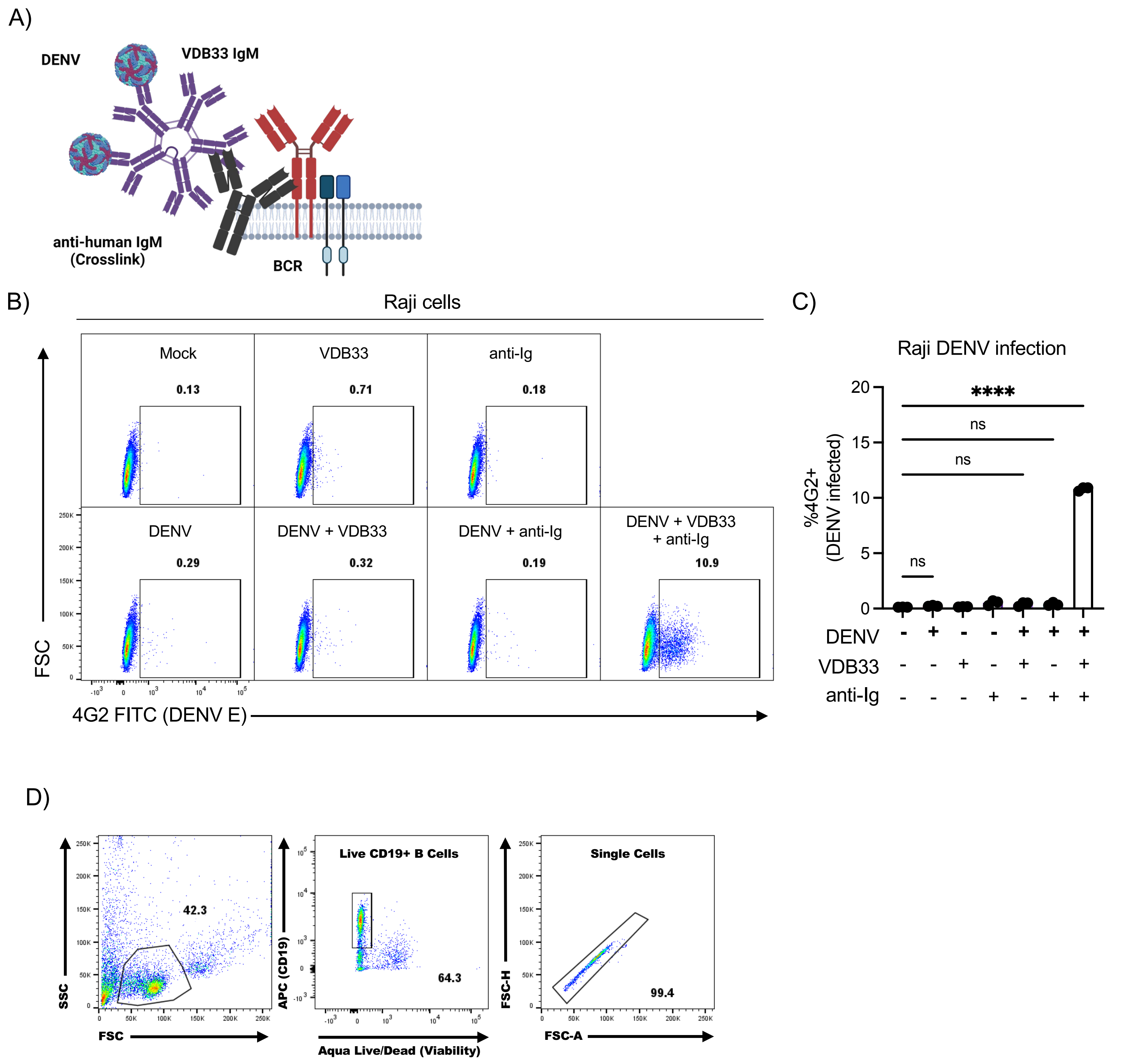


**Supplemental Figure 6. A)** Schematic representation of DENV/Ig/BCR crosslinking assay. **B**) Representative flow cytometry plot showing the frequency of DENV-4 infected Raji cells under the indicated culture conditions. Detection of DENV-infected cells was performed by staining fixed/permeabilized cells with a FITC-conjugated 4G2 antibody. **C**) Quantification of DENV-4 infected Raji cells under the indicated culture conditions. **D)** Gating scheme for B cell infection analysis utilizing the DENV/Ig/BCR crosslinking assay.

Supplemental Figure 7

A)

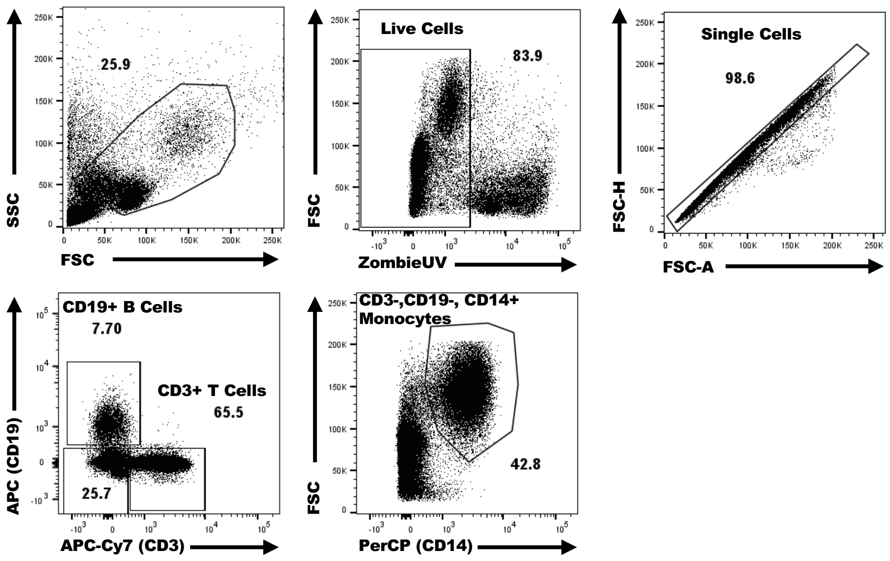


**Supplemental Figure 7.** Gating scheme for DHIM3 PBMC analysis
