## Supplemental Tables for "B cell receptor dependent enhancement of dengue virus infection"

**Supplemental Table 1.** Quality control metrics for scRNAseq data.

| **Sample** | **Number of reads** | **Estimated number of cells** | **Mean reads per cell** | **Median genes per cell** | **Median UMI counts per cell** |
| --- | --- | --- | --- | --- | --- |
| **Control** | 239 120 923 | 3 419 | 69 939 | 4 004 | 17 248 |
| **DENV-2** | 293 412 699 | 3 274 | 89 619 | 3 933 | 17 115 |

**Supplemental Table 2.** Infected cell characteristics across culture conditions.

| **Sample** | Total cells  n | (+) RNA only  n (%) | (-) RNA only  n (%) | (+) and (-) RNA  n (%) | **Percentage productively infected cells (%)** |
| --- | --- | --- | --- | --- | --- |
| **Control** | 3089 | 4 (0) | 0 (0) | 3 (0) | **0** |
| **DENV** | 2874 | 323 (11.2) | 46 (1.6) | 158 (5.5) | **5.5** |

**Supplemental Table 3.** Differentially expressed genes, DENV-2 infected cells and uninfected mock cells

| Gene | p_val | avg_log2FC | pct.1 | pct.2 | p_val_adj |
| --- | --- | --- | --- | --- | --- |
| DENV2 | 0 | 7.78946458 | 1 | 0 | 0 |
| DENV2neg | 0 | 5.64984623 | 1 | 0 | 0 |
| RPS29 | 7.25999011232681e-83 | 1.35628816 | 0.981 | 0.986 | 1.46564680387654e-78 |
| RPS21 | 1.4044602476356e-57 | 0.57383511 | 0.987 | 0.993 | 2.83532434792676e-53 |
| MT-ATP8 | 3.02340174175545e-54 | 1.01227813 | 0.994 | 0.97 | 6.10364343625591e-50 |
| RPL37A | 1.16825240324941e-53 | 0.53082581 | 0.981 | 0.994 | 2.35846795167992e-49 |
| ATP5ME | 1.74628912311265e-53 | 0.88281685 | 0.962 | 0.914 | 3.52540848173981e-49 |
| RPS27 | 1.52768057031352e-48 | 0.56638913 | 0.994 | 0.993 | 3.08408153534894e-44 |
| MT-ND3 | 1.39160781456075e-47 | 0.73113374 | 0.994 | 0.987 | 2.80937785603525e-43 |
| RPL38 | 1.39618780958835e-46 | 0.61665388 | 0.994 | 0.983 | 2.81862394999695e-42 |
| RPL37 | 2.10120092693786e-46 | 0.46669258 | 0.987 | 0.993 | 4.24190443130216e-42 |
| RMRP | 3.98916886228975e-46 | 0.86657363 | 0.728 | 0.251 | 8.05333409919055e-42 |
| NDUFB1 | 4.39891030636231e-38 | 0.71358662 | 0.949 | 0.863 | 8.88052012648424e-34 |
| RPL26 | 1.08898352981008e-34 | -0.5123385 | 0.968 | 0.99 | 2.19843994998059e-30 |
| UBD | 5.77634064960661e-33 | 0.34065402 | 0.487 | 0.141 | 1.16612765034258e-28 |
| ATP5F1E | 9.27021771196101e-33 | 0.45398847 | 0.968 | 0.988 | 1.87147155169069e-28 |
| RPS15A | 2.4407716955149e-28 | -0.4029856 | 0.987 | 0.994 | 4.92742989890547e-24 |
| NME2 | 1.06047926068114e-27 | 0.52578956 | 0.949 | 0.863 | 2.14089553146308e-23 |
| OAZ1 | 1.56817937998613e-24 | -0.4862034 | 0.981 | 0.985 | 3.165840532316e-20 |
| RPS28 | 1.27749003607066e-22 | 0.33160919 | 0.994 | 0.994 | 2.57899688481945e-18 |
| COX16 | 3.40763827400769e-22 | 0.46417728 | 0.867 | 0.654 | 6.87934014756672e-18 |
| ROMO1 | 9.06172504820304e-22 | 0.46921768 | 0.949 | 0.945 | 1.82938105273123e-17 |
| RPL36 | 1.61912291452079e-21 | 0.29645018 | 0.994 | 0.992 | 3.26868533983457e-17 |
| MIF | 2.06282658931158e-21 | 0.42464314 | 0.975 | 0.984 | 4.16443431850221e-17 |
| ALDOA | 2.80330944601422e-21 | 0.48325969 | 0.975 | 0.937 | 5.6593211096135e-17 |
| EIF5B | 5.69182820695662e-21 | 0.51527792 | 0.956 | 0.89 | 1.1490662784204e-16 |
| RPL39 | 1.11630036014419e-20 | 0.36510245 | 0.981 | 0.992 | 2.25358716705909e-16 |
| NDUFA3 | 3.76229184241984e-20 | 0.45464489 | 0.924 | 0.81 | 7.59531477147716e-16 |
| PTPRCAP | 4.82252370572321e-20 | 0.26526699 | 0.62 | 0.285 | 9.73571085711401e-16 |
| MTRNR2L12 | 1.60868694252421e-19 | -0.6386891 | 0.994 | 0.992 | 3.24761719956788e-15 |
| YWHAE | 1.29109630290322e-18 | 0.38938592 | 0.975 | 0.97 | 2.60646521630102e-14 |
| RPSA | 2.78908460063567e-18 | -0.2891266 | 0.987 | 0.989 | 5.63060399176328e-14 |
| DDT | 1.49190884199643e-17 | -0.4452174 | 0.873 | 0.929 | 3.01186557022239e-13 |
| ARPC1B | 2.75616116535174e-17 | 0.36627286 | 0.873 | 0.658 | 5.56413816061209e-13 |
| NEDD8 | 1.15235661387404e-16 | -0.4267738 | 0.88 | 0.928 | 2.32637753208891e-12 |
| RPS3A | 2.22088635006074e-16 | -0.2973643 | 0.987 | 0.996 | 4.48352536350263e-12 |
| TMA7 | 1.0108239307923e-15 | 0.33225826 | 0.994 | 0.985 | 2.0406513514835e-11 |
| EIF3J | 9.70854194640473e-15 | 0.38694512 | 0.861 | 0.659 | 1.95996044814019e-10 |
| RPL34 | 2.71187397446657e-14 | 0.28326956 | 0.987 | 0.994 | 5.47473117965312e-10 |
| TOMM5 | 3.61786651933212e-14 | 0.43725135 | 0.943 | 0.874 | 7.30374892922769e-10 |
| PSMA6 | 6.41696007303664e-14 | 0.37102156 | 0.911 | 0.757 | 1.29545589954464e-09 |
| ATP5MD | 1.00706280856607e-13 | 0.33729591 | 0.968 | 0.957 | 2.03305839793318e-09 |
| SLIRP | 1.2755346583616e-13 | 0.42725472 | 0.956 | 0.93 | 2.5750493683004e-09 |
| CMSS1 | 2.74354398831438e-13 | 0.25976057 | 0.677 | 0.429 | 5.53866660360908e-09 |
| PET100 | 7.84949360704804e-13 | 0.3116727 | 0.88 | 0.712 | 1.58465576939086e-08 |
| RUNX3 | 7.85331975575218e-13 | 0.37087152 | 0.886 | 0.721 | 1.58542819229125e-08 |
| DDX1 | 8.27141834192928e-13 | 0.34850161 | 0.823 | 0.646 | 1.66983393486868e-08 |
| MIR155HG | 1.47645514321639e-12 | 0.53260008 | 0.797 | 0.545 | 2.98066764312524e-08 |
| TNFAIP2 | 1.7574352966517e-12 | 0.60251258 | 0.937 | 0.82 | 3.54791037688044e-08 |
| ABCF1 | 3.57749450564282e-12 | 0.37302376 | 0.899 | 0.75 | 7.22224590799173e-08 |
| SAMSN1 | 5.97179491289947e-12 | 0.43817783 | 0.804 | 0.575 | 1.20558595701614e-07 |
| NDUFA11 | 7.30134428300667e-12 | -0.3284874 | 0.785 | 0.842 | 1.47399538385339e-07 |
| PA2G4 | 8.73064851011436e-12 | 0.44609962 | 0.949 | 0.928 | 1.76254332122189e-07 |
| METAP2 | 1.25492811923672e-11 | 0.3386349 | 0.911 | 0.811 | 2.53344888711509e-07 |
| EIF2S2 | 2.25054359597641e-11 | 0.33105807 | 0.943 | 0.91 | 4.54339741155719e-07 |
| COX6A1 | 2.27161015403552e-11 | -0.2942879 | 0.981 | 0.982 | 4.58592657896692e-07 |
| NARS | 2.62334462180443e-11 | 0.26499175 | 0.829 | 0.606 | 5.29600812249878e-07 |
| PPA1 | 2.71720292700571e-11 | 0.30753662 | 0.968 | 0.959 | 5.48548926903913e-07 |
| PPDPF | 4.02509472606887e-11 | -0.3969425 | 0.924 | 0.945 | 8.12586123298784e-07 |
| RPS4X | 5.87743488249973e-11 | -0.2558316 | 0.994 | 0.995 | 1.18653655407905e-06 |
| S100A4 | 7.41867516922651e-11 | 0.51334713 | 0.772 | 0.577 | 1.49768214316345e-06 |
| H3F3A | 8.32096806568795e-11 | -0.2919772 | 0.975 | 0.991 | 1.67983703310108e-06 |
| MRPL23 | 1.16849639390254e-10 | -0.2976701 | 0.405 | 0.586 | 2.35896052001044e-06 |
| TAF11 | 1.32539947611917e-10 | 0.25587334 | 0.759 | 0.526 | 2.67571646238939e-06 |
| PPP2CA | 1.3578764262815e-10 | -0.3642489 | 0.658 | 0.768 | 2.7412809293771e-06 |
| H3F3B | 1.84321826352206e-10 | -0.3096507 | 0.987 | 0.993 | 3.72108903039833e-06 |
| MT-ATP6 | 2.7545535464703e-10 | -0.32063 | 1 | 0.994 | 5.56089269961423e-06 |
| DENR | 6.15790294291304e-10 | 0.27764285 | 0.886 | 0.756 | 1.24315744611528e-05 |
| SLC25A5 | 9.78785086703137e-10 | -0.3025437 | 0.943 | 0.979 | 1.97597133303629e-05 |
| NDUFA1 | 1.21867464662305e-09 | 0.27854151 | 0.968 | 0.941 | 2.46026037660262e-05 |
| MT-ND2 | 1.36006841469263e-09 | 0.30662285 | 1 | 0.985 | 2.74570611558149e-05 |
| AL138963.3 | 2.63463105579513e-09 | -0.5445581 | 0.741 | 0.804 | 5.31879317543921e-05 |
| CCR7 | 3.42496449185524e-09 | 0.70978491 | 0.354 | 0.172 | 6.91431831615736e-05 |
| SRSF9 | 3.52210730495601e-09 | -0.2780663 | 0.968 | 0.966 | 7.11043022724518e-05 |
| CORO1A | 6.55693077567514e-09 | -0.3485382 | 0.949 | 0.958 | 0.00013237 |
| DPYSL2 | 8.88444384567165e-09 | 0.28215948 | 0.772 | 0.568 | 0.00017936 |
| SNRPB2 | 1.05693364179265e-08 | 0.30624269 | 0.892 | 0.797 | 0.00021337 |
| PRDX2 | 1.21359193980804e-08 | -0.3082327 | 0.949 | 0.966 | 0.000245 |
| ERH | 2.13954803762635e-08 | -0.2748098 | 0.962 | 0.978 | 0.00043193 |
| IGLC3 | 3.01192707291274e-08 | 0.40509046 | 0.93 | 0.752 | 0.00060805 |
| FNBP1 | 3.04446929763137e-08 | 0.29256032 | 0.905 | 0.813 | 0.00061462 |
| NAPSA | 4.76436719908735e-08 | -0.3482562 | 0.538 | 0.639 | 0.00096183 |
| ARPC5 | 7.09231545556128e-08 | -0.2946798 | 0.968 | 0.979 | 0.0014318 |
| MKNK2 | 1.2468912670071e-07 | 0.3506819 | 0.861 | 0.722 | 0.00251722 |
| CD83 | 1.31044635033675e-07 | 0.41216608 | 0.804 | 0.647 | 0.00264553 |
| HLA-C | 1.34908219653009e-07 | -0.3426838 | 0.956 | 0.966 | 0.00272353 |
| DDX21 | 1.74955084302371e-07 | 0.33254476 | 0.918 | 0.9 | 0.00353199 |
| PPP1CA | 1.85579640066829e-07 | -0.2562549 | 0.937 | 0.949 | 0.00374648 |
| ATP6V1G1 | 1.88510744852717e-07 | -0.3312708 | 0.943 | 0.952 | 0.00380565 |
| GPR157 | 1.9375995077652e-07 | 0.27679599 | 0.437 | 0.261 | 0.00391163 |
| LCP1 | 2.25436168594684e-07 | 0.2871279 | 0.987 | 0.985 | 0.00455111 |
| MALAT1 | 2.26555259303704e-07 | 0.33674625 | 1 | 0.995 | 0.0045737 |
| SKP1 | 2.3382582128986e-07 | -0.2617133 | 0.943 | 0.95 | 0.00472048 |
| CCNI | 2.5862007400521e-07 | -0.274419 | 0.937 | 0.954 | 0.00522102 |
| IGKV3-15 | 2.64167659909777e-07 | -1.3226319 | 0.082 | 0.252 | 0.00533302 |
| PDCD11 | 2.80322090233665e-07 | 0.2694218 | 0.728 | 0.547 | 0.00565914 |
| CCNG1 | 3.13780313320918e-07 | -0.3096497 | 0.918 | 0.908 | 0.0063346 |
| WDR83OS | 3.33462018465226e-07 | -0.2857171 | 0.924 | 0.933 | 0.00673193 |
| TMEM14B | 4.34447239092933e-07 | -0.2755085 | 0.842 | 0.871 | 0.00877062 |
| IGKV3D-11 | 5.36806135548254e-07 | -1.1740944 | 0.082 | 0.25 | 0.01083704 |
| DYNLL1 | 5.37537015403036e-07 | -0.3371743 | 0.93 | 0.964 | 0.0108518 |
| LIMD2 | 5.69800488403979e-07 | -0.2836384 | 0.975 | 0.963 | 0.01150313 |
| ANP32B | 6.29284332714697e-07 | -0.2804014 | 0.968 | 0.968 | 0.01270399 |
| IGKV3-11 | 7.62445584680517e-07 | -1.5906087 | 0.095 | 0.256 | 0.01539225 |
| TNNI1 | 8.02541581968897e-07 | 0.3400626 | 0.519 | 0.317 | 0.01620171 |
| PLEC | 8.52500839120461e-07 | 0.35477235 | 0.753 | 0.591 | 0.01721029 |
| PSMD9 | 9.01146687812747e-07 | -0.2714013 | 0.582 | 0.69 | 0.01819235 |
| EIF3H | 9.90101493396531e-07 | -0.2639272 | 0.956 | 0.967 | 0.01998817 |
| CD44 | 9.91814559212962e-07 | 0.38893039 | 0.241 | 0.114 | 0.02002275 |
| IGKC | 1.15333776199585e-06 | -1.2112693 | 0.114 | 0.276 | 0.02328358 |
| SCD | 1.87330420715242e-06 | 0.26534152 | 0.854 | 0.761 | 0.03781827 |
| CCL17 | 1.88548054439542e-06 | 0.59604529 | 0.19 | 0.08 | 0.03806408 |
| FTL | 1.90599002852873e-06 | -0.3101528 | 0.981 | 0.994 | 0.03847813 |
| IGHV5-51 | 1.99311771255404e-06 | -1.49403 | 0.095 | 0.254 | 0.04023706 |
| FLNA | 2.02226636035301e-06 | 0.31834453 | 0.911 | 0.87 | 0.04082551 |
| TIMP1 | 2.02410921494935e-06 | 0.30435698 | 0.924 | 0.851 | 0.04086272 |
| CCL22 | 2.14564901093193e-06 | 0.60350096 | 0.532 | 0.345 | 0.04331636 |
| HYOU1 | 2.15838041353401e-06 | 0.25599756 | 0.829 | 0.695 | 0.04357338 |

**Supplemental Table 4.** Differentially expressed pathways between DENV-2 infected 7B9 cells and uninfected mock cells

| **Ingenuity Canonical Pathways** | **-log(p-value)** | **Ratio** | **z-score** |
| --- | --- | --- | --- |
| Eukaryotic Translation Initiation | 23.3 | 0.156 | 1.606 |
| Response of EIF2AK4 (GCN2) to amino acid deficiency | 19.6 | 0.155 | 1.5 |
| EIF2 Signaling | 19.3 | 0.087 | 1.667 |
| Nonsense-Mediated Decay (NMD) | 18.7 | 0.137 | 1 |
| Eukaryotic Translation Termination | 18.6 | 0.16 | 1.291 |
| Eukaryotic Translation Elongation | 18.5 | 0.158 | 1.291 |
| Selenoamino acid metabolism | 17.7 | 0.14 | 1.291 |
| SRP-dependent cotranslational protein targeting to membrane | 17.2 | 0.13 | 1.291 |
| Major pathway of rRNA processing in the nucleolus and cytosol | 16.8 | 0.0914 | 1.698 |
| Electron transport, ATP synthesis, and heat production by uncoupling proteins | 12.1 | 0.0938 | 1.732 |
| Regulation of eIF4 and p70S6K Signaling | 10.2 | 0.0652 | #NUM! |
| Oxidative Phosphorylation | 9.92 | 0.0893 | 1.265 |
| mTOR Signaling | 9.44 | 0.0561 | #NUM! |
| Sirtuin Signaling Pathway | 6.92 | 0.0378 | -1.89 |
| Coronavirus Pathogenesis Pathway | 6.31 | 0.0441 | -0.333 |
| Granzyme A Signaling | 5.89 | 0.0811 | -1.633 |
| Neutrophil Extracellular Trap Signaling Pathway | 5.56 | 0.0275 | 0.905 |
| Mitochondrial Dysfunction | 5.31 | 0.0291 | -1.265 |
| Cristae formation | 4.9 | 0.129 | 1 |
| Binding and Uptake of Ligands by Scavenger Receptors | 4.85 | 0.0536 | -0.816 |
| rRNA processing | 4.84 | 0.125 | #NUM! |
| Signaling by the B Cell Receptor (BCR) | 4.82 | 0.0412 | -1.134 |
| Hematoma Resolution Signaling Pathway | 4.54 | 0.031 | 1.414 |
| Fc epsilon receptor (FCERI) signaling | 4.29 | 0.034 | -1.134 |
| HIPPO signaling | 4.26 | 0.0575 | #NUM! |
| C-type lectin receptors (CLRs) | 4.21 | 0.0414 | 0 |
| Fcgamma receptor (FCGR) dependent phagocytosis | 4.02 | 0.0382 | -0.816 |
| Parkinson's Signaling Pathway | 4.01 | 0.0261 | -1.414 |
| Eumelanin Biosynthesis | 3.89 | 0.5 | #NUM! |
| Iron uptake and transport | 3.81 | 0.069 | -2 |
| NIK-->noncanonical NF-kB signaling | 3.75 | 0.0667 | -1 |
| Mitotic G2-G2/M phases | 3.46 | 0.0302 | -0.816 |
| Neutrophil degranulation | 3.41 | 0.0189 | 0.333 |
| Glucocorticoid Receptor Signaling | 3.4 | 0.0172 | #NUM! |
| Cell surface interactions at the vascular wall | 3.33 | 0.0284 | 0 |
| TP53 Regulates Metabolic Genes | 3.11 | 0.0455 | 0 |
| Degradation of beta-catenin by the destruction complex | 3.04 | 0.0435 | -1 |
| ABC-family proteins mediated transport | 2.89 | 0.0396 | 1 |
| Cell Cycle Checkpoints | 2.76 | 0.0221 | 0 |
| Macrophage Alternative Activation Signaling Pathway | 2.7 | 0.0263 | 1.342 |
| FAT10 Signaling Pathway | 2.62 | 0.0526 | #NUM! |
| Immunoregulatory interactions between a Lymphoid and a non-Lymphoid cell | 2.59 | 0.0248 | -1.342 |
| Metabolism of polyamines | 2.58 | 0.0508 | #NUM! |
| Estrogen Receptor Signaling | 2.51 | 0.0171 | #NUM! |
| Mitotic G1 phase and G1/S transition | 2.48 | 0.0305 | #NUM! |
| Complement cascade | 2.43 | 0.0296 | #NUM! |
| Regulation of RUNX2 expression and activity | 2.32 | 0.0411 | #NUM! |
| Glioma Invasiveness Signaling | 2.32 | 0.0411 | #NUM! |
| Mitotic Metaphase and Anaphase | 2.31 | 0.0213 | -0.447 |
| p70S6K Signaling | 2.24 | 0.0138 | #NUM! |
| Neddylation | 2.23 | 0.0203 | -0.447 |
| Epithelial Adherens Junction Signaling | 2.19 | 0.0253 | 1 |
| Signaling by NOTCH4 | 2.16 | 0.0361 | #NUM! |
| Inhibition of ARE-Mediated mRNA Degradation Pathway | 2.15 | 0.0245 | #NUM! |
| Phagosome Maturation | 2.14 | 0.0244 | #NUM! |
| Cyclins and Cell Cycle Regulation | 2.12 | 0.0349 | #NUM! |
| Regulation of mitotic cell cycle | 2.09 | 0.0341 | #NUM! |
| KEAP1-NFE2L2 pathway | 2.05 | 0.033 | #NUM! |
| Crosstalk between Dendritic Cells and Natural Killer Cells | 2.05 | 0.033 | #NUM! |
| Protein Ubiquitination Pathway | 2.04 | 0.0183 | -0.447 |
| Actin Nucleation by ARP-WASP Complex | 2.03 | 0.0323 | #NUM! |
| Transcriptional regulation by RUNX3 | 1.99 | 0.0312 | #NUM! |
| S Phase | 1.94 | 0.03 | #NUM! |
| RHO GTPases Activate WASPs and WAVEs | 1.91 | 0.0556 | #NUM! |
| Protein Kinase A Signaling | 1.9 | 0.0146 | #NUM! |
| Gene and protein expression by JAK-STAT signaling after IL-12 stimulation | 1.89 | 0.0541 | #NUM! |
| Pyrophosphate hydrolysis | 1.86 | 0.333 | #NUM! |
| TCF dependent signaling in response to WNT | 1.85 | 0.0201 | 0 |
| Regulation of TP53 Expression and Degradation | 1.85 | 0.0513 | #NUM! |
| Mitotic Prometaphase | 1.82 | 0.0197 | 0 |
| Hedgehog 'off' state | 1.8 | 0.0265 | #NUM! |
| Regulation of Actin-based Motility by Rho | 1.78 | 0.0261 | #NUM! |
| Synthesis of DNA | 1.74 | 0.0252 | #NUM! |
| ERK/MAPK Signaling | 1.74 | 0.0186 | 1 |
| Interleukin-10 signaling | 1.73 | 0.0444 | #NUM! |
| RHOGDI Signaling | 1.71 | 0.0182 | -1 |
| TCR signaling | 1.67 | 0.0238 | #NUM! |
| Interleukin-1 family signaling | 1.65 | 0.0233 | #NUM! |
| Cell Cycle: G2/M DNA Damage Checkpoint Regulation | 1.63 | 0.0392 | #NUM! |
| Response to elevated platelet cytosolic Ca2+ | 1.62 | 0.0227 | #NUM! |
| Regulation of Apoptosis | 1.6 | 0.0377 | #NUM! |
| Signaling by TGF-beta Receptor Complex | 1.6 | 0.0377 | #NUM! |
| RAC Signaling | 1.58 | 0.0219 | #NUM! |
| B Cell Development | 1.57 | 0.0123 | #NUM! |
| Iron homeostasis signaling pathway | 1.57 | 0.0217 | #NUM! |
| Intrinsic Pathway for Apoptosis | 1.57 | 0.0364 | #NUM! |
| RHO GTPases Activate Formins | 1.56 | 0.0216 | #NUM! |
| Communication between Innate and Adaptive Immune Cells | 1.52 | 0.00966 | -1 |
| Circadian Clock | 1.51 | 0.0339 | #NUM! |
| Semaphorin Signaling in Neurons | 1.48 | 0.0328 | #NUM! |
| Primary Immunodeficiency Signaling | 1.48 | 0.0328 | #NUM! |
| Cytoprotection by HMOX1 | 1.47 | 0.0323 | #NUM! |
| UFMylation Signaling Pathway | 1.47 | 0.0323 | #NUM! |
| IL-15 Signaling | 1.44 | 0.0114 | #NUM! |
| Hedgehog ligand biogenesis | 1.43 | 0.0308 | #NUM! |
| FcγRIIB Signaling in B Lymphocytes | 1.42 | 0.0113 | #NUM! |
| TNFR2 non-canonical NF-kB pathway | 1.4 | 0.0294 | #NUM! |
| Remodeling of Epithelial Adherens Junctions | 1.4 | 0.0294 | #NUM! |
| OAS antiviral response | 1.39 | 0.111 | #NUM! |
| Sucrose Degradation V (Mammalian) | 1.39 | 0.111 | #NUM! |
| Cell Cycle: G1/S Checkpoint Regulation | 1.39 | 0.029 | #NUM! |
| Huntington's Disease Signaling | 1.36 | 0.0141 | #NUM! |
| Netrin Signaling | 1.33 | 0.0173 | #NUM! |
| Cellular response to hypoxia | 1.32 | 0.0267 | #NUM! |
| Caveolar-mediated Endocytosis Signaling | 1.32 | 0.0267 | #NUM! |
| WNT/β-catenin Signaling | 1.32 | 0.0172 | #NUM! |

**Supplemental Table 5.** Reagents for flow cytometry analysis

| **Antibody** | **Clone** | **Dilution** | **Catalog #** | **Lot #** |
| --- | --- | --- | --- | --- |
| Anti-human IgA AF647 | N/A | 1:200 | Southern Biotech, 2050-31 | G0919-V490B |
| Anti-human IgG AF647 | N/A | 1:200 | Southern Biotech, 2040-31 | B3919-M950B |
| Anti-human IgM AF647 | N/A | 1:200 | Southern Biotech, 2020-31 | D2219-R121 |
| Anti-human CD19 APC | HIB19 | 1:200 | Biolegend, 302212 | B386154 |
| Anti-human CD14 PerCP | M5E2 | 1:50 | Biolegend, 301848 | B371416 |
| Anti-human CD3 APC-Cy7 | OKT3 | 1:200 | Biolegend, 317342 | B367892 |
| Anti-human DC-SIGN | 9E9A8 | 1:200 | Biolegend, 330112 | B386835 |
| Anti-human PDGFRα | 16A1 | 1:200 | Biolegend, 323512 | B388290 |
| ZombieUV | N/A | 1:200 | Biolegend, 77474 | B385840 |
| Aqua Live/Dead | N/A | 1:500 | Invitrogen, L34957 | 2204201 |
| DENV Envelop (4G2) FITC | N/A | 4ug/mL | Envigo Bioproducts, Inc, CON004 | N/A |
| Anti-mouse IgG Fab2 AF647 | N/A | 0.4ug/mL | Cell Signaling Technology, Inc, 4410S | 17 |
| Near-IR fluorescent reactive dye | N/A | 1:1000 | Invitrogen | L34975A |
| Anti-human CD19-PerCP-Cy5.5 | HIB19 | 1:50 | Biolegend, 302230 | B402462 |
| Anti-human IgG Fc APC | M1310G05 | 1:50 | Biolegend, 410712 | B405400 |
